# Capturing Multicellular System Designs Using the Synthetic Biology Open Language (SBOL)

**DOI:** 10.1101/463844

**Authors:** Bradley Brown, Christian Atallah, James Alastair McLaughlin, Göksel Misirli, Ángel Goñi-Moreno, Nicholas Roehner, David James Skelton, Bryan Bartley, Jacob Beal, Chueh Loo Poh, Irina Dana Ofiteru, Anil Wipat

## Abstract

Synthetic biology aims to improve the development of biological systems and increase their reproducibility through the use of engineering principles, such as standardisation and modularisation. It is important that these systems can be represented and shared in a standard way to ensure they are easily understood, reproduced, and utilised by other researchers. The Synthetic Biology Open Language (SBOL) is a data standard for sharing biological designs and information about their implementation and characterisation. Thus far, this standard has been used to represent designs in homogeneous systems, where the same design is implemented in every cell. In recent years there has been increasing interest in multicellular systems, where biological designs are split across multiple cells to optimise the system behaviour and function. Here we show how the SBOL standard can be used to represent such multicellular systems, and hence how researchers can better share designs with the community.

## 1 Introduction

The increasing popularity of synthetic biology has yielded a wealth of biological systems which have been designed, implemented, and characterised (*1, 2*). These systems span a wide range of functionalities, from fundamental genetic circuits such as oscillators (*3*) and toggle switches (*4*), to applied devices such as biosensors (*5*) and microbial factories (*6*). Classically, these biological devices have been designed and implemented as homogeneous systems, where identical genetic circuits are intended to be expressed by every cell in the population. Whilst this approach has yielded promising results, the complexity and optimisation of such systems can become limited for certain applications (*7*). One reason for these limitations is that the larger genetic circuits required by more complex systems place cells under metabolic strain, resulting in sub-optimal performance. Another reason is that elements in large and complex designs tend to be drawn from many diverse sources, which have different optimal environments (*8*). Therefore, the host cell chosen to express these circuits tends to be a compromise which limits overall optimal performance.

In recent years, there has been a new wave of engineered biological designs using multi-cellular systems, where different elements of the system are split and expressed by separate cells. These cells can be engineered to communicate with each other to create a co-culture of cells which, together, can perform a desired function (*9–13*). This approach can reduce the metabolic burden on individual cells, which now only have to perform a fragment of the overall system, and optimal host cells can be chosen for each element of the design.

One of the major hallmarks of synthetic biology is standardisation, which aims to increase the reproducibility of engineered biological systems and facilitate re-use by other researchers. To fully realise this aim, it is important that information about the design, implementation, and characterisation of engineered systems can be easily shared with and understood by other members of the synthetic biology community. The Synthetic Biology Open Language (SBOL) (*14*) has been developed by a community of synthetic biologists to capture information about engineered systems in a standardised format. In a similar fashion to the way that file formats such as the GenBank flat file format (GBF) were developed to capture information about natural biological systems (*15*), SBOL enables the design-build-test cycle to be standardised by storing information required at each stage. This information (designs, build plans, implementation details, and experimental information/results) aids the sharing of information between researchers and labs (*16, 17*). Thus far, SBOL has been used to represent designs implemented in homogeneous cultures as opposed to multicellular systems. Here, we discuss how the SBOL standard (version 2.2.1(*18*)) could be used to represent multicellular system designs. Unified Modeling Language (UML) diagrams and examples are used throughout to illustrate how different aspects of a system’s design can be captured. Additionally, it is discussed how a standard for representing multicellular systems can be used to assist synthetic biology researchers. It is intended that this paper will provide a basis for further discussions and refinements within the synthetic biology scientific community.

## 2 Methods

In this section, the relevant sections of the SBOL data model are explained. Additionally, the best practices and data model extensions proposed here to represent multicellular systems in SBOL are described in detail. The current version of SBOL at the time of writing is version 2.2.1(*18*).

### 2.1 The *ComponentDefinition* and *ModuleDefinition* Classes

The current SBOL data model has two main classes used to capture biological designs: *ComponentDefinition* and *ModuleDefinition. ComponentDefinition* is usually used to capture physical structures, such as DNA and proteins, whereas *ModuleDefinition* is used to group together biological entities in a design and define molecular interactions. The designs captured can range in complexity, from single parts (e.g. a promoter or protein), to devices composed of multiple parts (e.g. an expression construct). In the case of devices, each individual part must be described by a separate *ComponentDefinition* or *ModuleDefinition*. An instance of that part must then be created within the class describing the device using a *FunctionalComponent* class.

Instances of the *ModuleDefinition* class may contain interactions between biological entities in the design (e.g. a coding sequence CDS encodes a protein which represses a promoter), whereas instances of *ComponentDefinition* may not. Additionally, *ComponentDefinition* has both type and role properties, but *ModuleDefinition* has only role properties. The type property in SBOL is used to describe the category within which a biological entity falls (e.g. DNA molecule, small molecule, protein), and the ‘role’ property describes the intended function for an entity or design (e.g. metabolic pathway, AND gate, protein expression).

### 2.2 Ontologies in SBOL

An ontology can be thought of as a set of formal definitions for specific terms and relationships within a discipline. In synthetic biology, this allows a standardised language to be used when describing biological systems. A number of separate ontologies are used in SBOL to better describe entities within a system, and how those entities interact. For example, the ‘role’ property used by instances of the *ComponentDefinition* class can be defined by terms taken from the Sequence Ontology (SO) (*19*). Examples of terms used in this property are ‘Promoter’, ‘RBS’, and ‘CDS’. Other ontologies commonly used in SBOL include the Systems Biology Ontology (SBO) (*20*), the Gene Ontology (GO) (*21, 22*), and Chemical Entities of Biological Interest (CHEBI) (*23*).

### 2.3 Representing Cells in SBOL

Two approaches for representing cells in a biological system are proposed here. Both approaches are capable of capturing the same information: (i) taxonomy, (ii) interactions occurring within the cell, and (iii) components inside the cell (e.g. DNA molecules).

The first approach uses an instance of the *ModuleDefinition* class to represent the cell used in the design. This class instance has a role of ‘physical compartment’ from the Synthetic Biology Ontology (SBO:0000290). The species and strain of the cell is captured by annotating the *ModuleDefinition* class instance with an ‘organism’ property, which is an URI (Uniform Resource Identifier) for a relevant entry in the NCBI Taxonomy Database. In the case of organisms not currently in the NCBI database, a URI pointing to another database or a description of the organism can be used. Entities relevant to the biological design, and which can be found inside the cell, are captured using instances of the *FunctionalComponent* class contained by the *ModuleDefinition*. Instances of the *Interaction* and *Participation* classes are used to capture interactions between these entities.

The second approach uses an instance of the *ModuleDefinition* class to represent a system which contains the cell. The cell is represented by an instance of the *FunctionalComponent* class inside the *ModuleDefinition*. A separate *ComponentDefinition* instance is used to capture information about the species and strain of the cell in the design. This *ComponentDefinition* has a ‘type’ of ‘cell’ from the Gene Ontology (GO:0005623), and a role of ‘physical compartment’ (SBO:0000290). Taxonomic information is captured by annotating the class instance with a URI for an entry in the NCBI Taxonomy Database. The *ComponentDefinition* instance is used as the value of the ‘definition’ property of the *FunctionalComponent* which represents the cell in the design to specify information about its taxonomy. Other entities which are relevant to this aspect of the design are also captured using instances of the *FunctionalComponent* class. Interactions which occur in this system are captured using the *Interaction* and *Participation* classes. Interactions which occur within the cell are specified by *Interaction* classes which contain the *FunctionalComponent* instance representing the cell as a participant with a ‘role’ of ‘physical compartment’.

For both approaches, an additional *Interaction* class instance could be used to define entities which are only present within the cell, i.e. they are not available to the rest of the system. This interaction has a ‘type’ of ‘containment’ (SBO:0000469), and has at least two participants. One of these participants is the cell, which has a ‘role’ of ‘physical compartment’, whilst the others are the contained entities, which have ‘roles’ of ‘contained’ (SBO:0000064).

### 2.4 Representing Designs with Multiple Cells

We are proposing three approaches for capturing designs which incorporate more than one cell. These approaches all utilise an instance of the *ModuleDefinition* class which has a ‘role’ of ‘functional compartment’. The first approach simply instantiates a *Module* class within this *ModuleDefinition* to represent each cell in the system. The *Module* classes are defined by an instance of the *ModuleDefinition* class which represents a cell, and is therefore compatible with either method of capturing cells described above. In this approach, interactions between cells are captured implicitly. Cells are defined as interacting if they contain or participate in interactions with the same entity, and are present in the same system. An abstract example of this is shown in Figure 2, where Cell 1 and Cell 2 both interact with Molecule A, and are incorporated in the same design.

**Figure 1:**
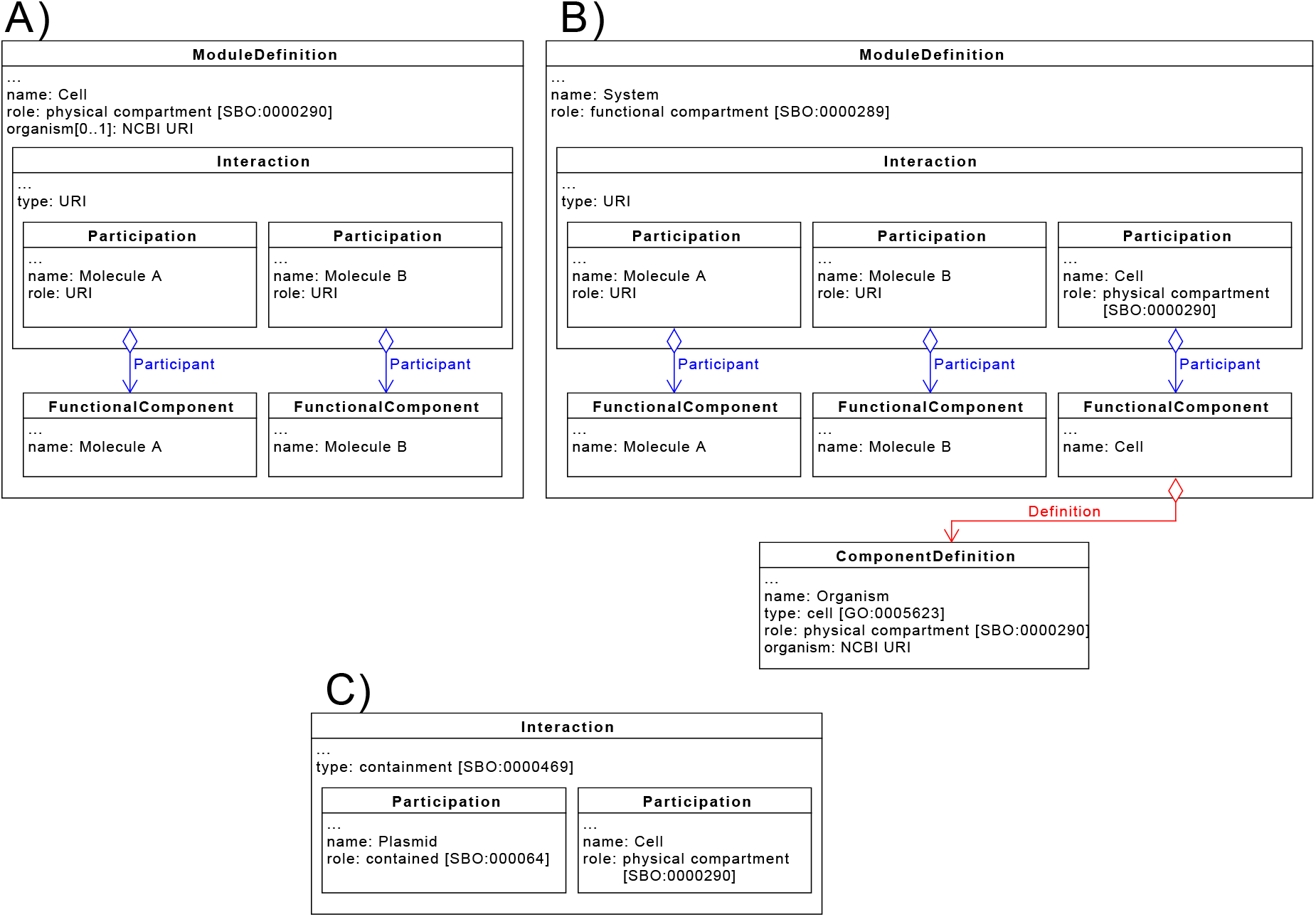
UML Diagram Depicting Representation of Cells in SBOL: The examples in A) and B) represent a cell which contains molecules ‘A’ and ‘B’, which interact in some way. A) First approach for capturing cell designs in SBOL. A *ModuleDefinition* is used to represent the entire cell. This *ModuleDefinition* has a role of ‘physical compartment’ from the Systems Biology Ontology (SBO) and is annotated with a URI pointing to an entry in the NCBI Taxonomy Database. Interactions occurring within the cell are specified using *Interaction* classes. B) Second approach for capturing cell designs in SBOL. A *ComponentDefinition* annotated with a URI pointing to an entry in the NCBI Taxonomy Database is used to capture information about the cell’s strain/species. The *ComponentDefinition* instance has a type of ‘Cell’ from the Gene Ontology (GO), and a role of ‘physical compartment’. An instance of the *ModuleDefinition* class is used to represent a system in which the cell is implemented. Entities, including the cell, are instantiated as *FunctionalComponents*, and process are captured using the *Interaction* class. Process which are contained within the cell are represented by including the cell as a participant with a role of ‘physical compartment’. C) Example of how the *Interaction* class can be used to explicitly confer that an entity is contained inside a cell. Here a plasmid is specified as being contained within a cell.

**Figure 2:**
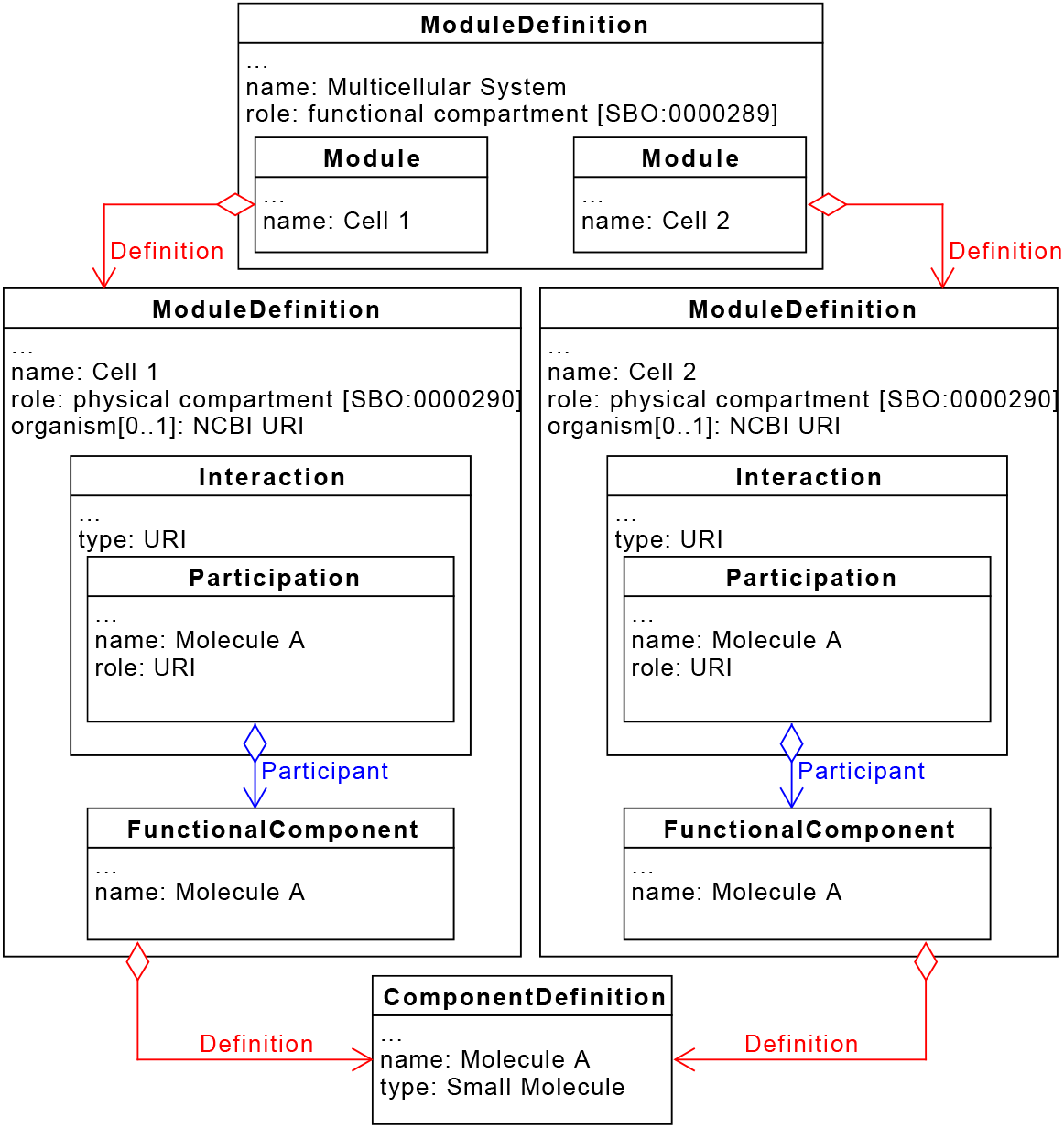
UML Diagram Depicting the First Suggestion for Representing Multicellular Designs in SBOL: Captured here is a design involving two cells which both interact with the small molecule ‘Molecule A’. Designs for Cell 1 and Cell 2 are captured using the approach depicted in *Figure 1A*, however this proposed approach is also compatible with the method shown in *Figure 1B*. The overall multicellular system is represented by a *ModuleDefinition* with a role of ‘functional compartment’, which is an SBO term. Both Cell 1 and Cell 2 are included in this design as an instance of the *Module* class. Interactions between the cells can be elucidated by comparing the entities which participate in processes defined within the cell designs. Cell 1 and Cell 2 in this design both have process which involve ‘Molecule A’, and hence can be deduced to have some form of intercellular interactions.

The second approach instantiates cells as a *FunctionalComponent* within the *ModuleDefinition* used to represent the multicellular system. Other entities present in the system, such as small molecules, are also instantiated as *FunctionalComponent* s. Instances of the *Interaction* classes are used to define interactions between the cells and other entities. In these interactions, cells have a role of functional compartment, which is an SBO term (SBO:0000289), to convey that the intracellular process is occurring within that cell. If a non-cell entity is instantiated in more than one *Interaction* class, then any cells also involved in these interactions are determined to have intercellular interactions. This is demonstrated in Figure 3. In the current version of SBOL (2.2.1), *FunctionalComponent* classes may only be defined by *ComponentDefinition* s, and not *ModuleDefinition* s. Therefore, the first approach for capturing cells described above is not compatible with this method, as no *ComponentDefinition* classes are used to describe the cell. The second method for capturing information about cells in a biological design is compatible with this approach; the *ComponentDefinition* classes used to capture information about the organism are used to define the *FunctionalComponent* instances, as demonstrated in Figure 3.

**Figure 3:**
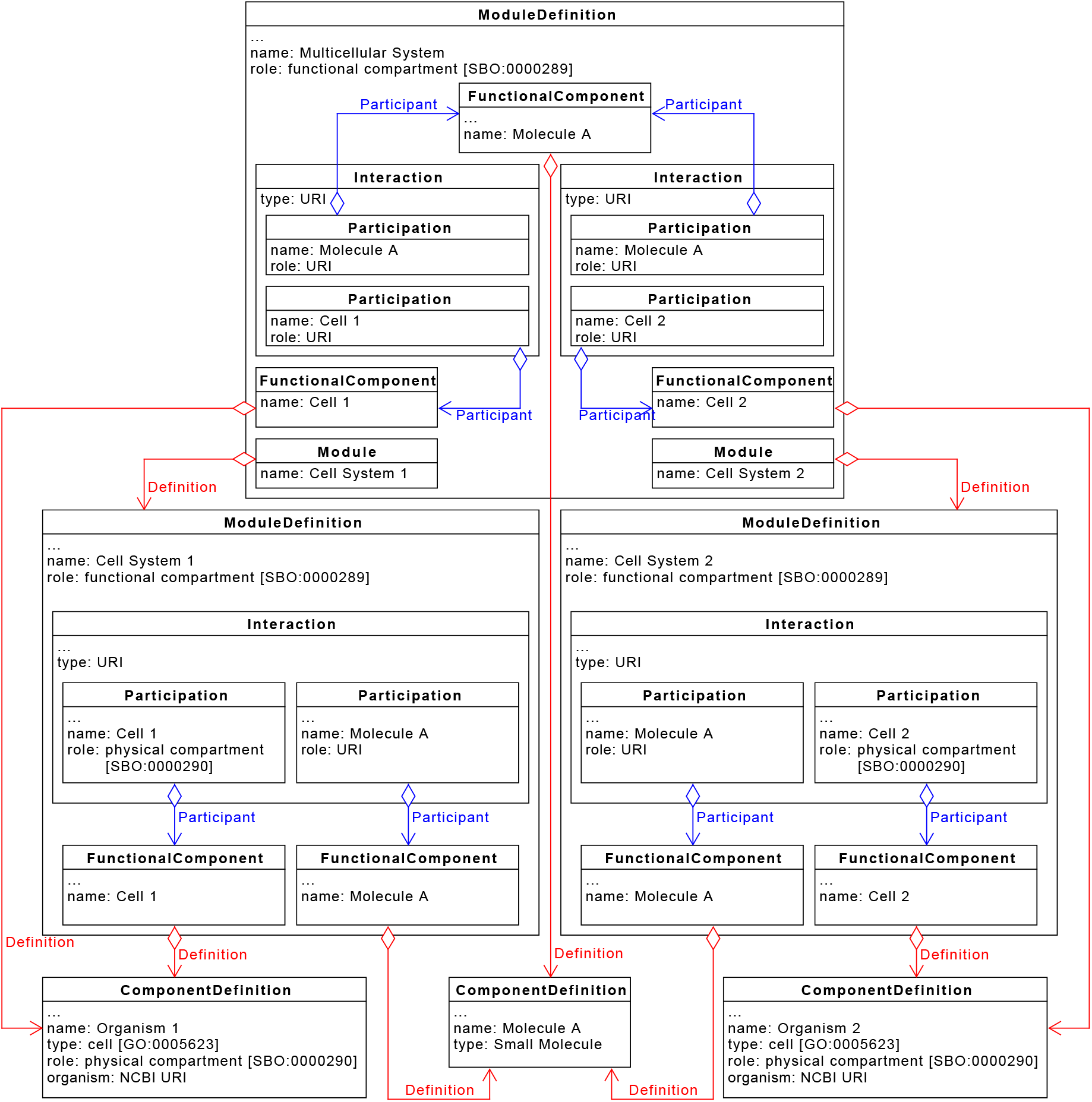
UML Diagram Depicting the Second Suggestion for Representing Multicellular Designs in SBOL: Captured here is a design involving two cells which both interact with the small molecule ‘Molecule A’. Designs for Cell 1 and Cell 2 are captured using the approach depicted in *Figure 1B*, as this proposed approach is not compatible with the method shown in *Figure 1A*. The overall multicellular system is represented by a *ModuleDefinition* with a role of ‘functional compartment’. Both Cell 1 and Cell 2 are included in this design as an instance of the *FunctionalComponent* class, which is defined by the *ComponentDefinition* instance which captures taxonomic information about the cell. Intercellular interactions are defined explicitly using the *Interaction* class. In this design, ‘Molecule A’ participates in processes within both Cell 1 and Cell 2.

The third approach proposed uses both *Module* and *FunctionalComponent* classes to represent cells, and any non-cell entities must also be instantiated as *FunctionalComponent* classes. It should be noted that this approach would not be compatible with the first method of representing cells as described above.

The *Module* instances are defined by the *ModuleDefinition* classes used to represent cells/cell systems as in the first approach. *FunctionalComponent* classes are also used to represent taxonomic information about the cell, and are defined by the *ComponentDefinition* classes which capture this information. Finally, the *Module* classes contain instances of the *MapsTo* class, which are used to explicitly capture links between the same entities present in multiple parts of the same design. Here, a *MapsTo* class with a ‘refinement’ value of ‘merge’ is used to link *FunctionalComponent* classes which represent a cell in the multicellular system to the *FunctionalComponent* class used to represent the same cell in the lower-level cell system design. Instances of the *MapsTo* class are also used to capture that non-cell entities in the multicellular system are identical to entities used in the cell system design. This is demonstrated in Figure 4.

**Figure 4:**
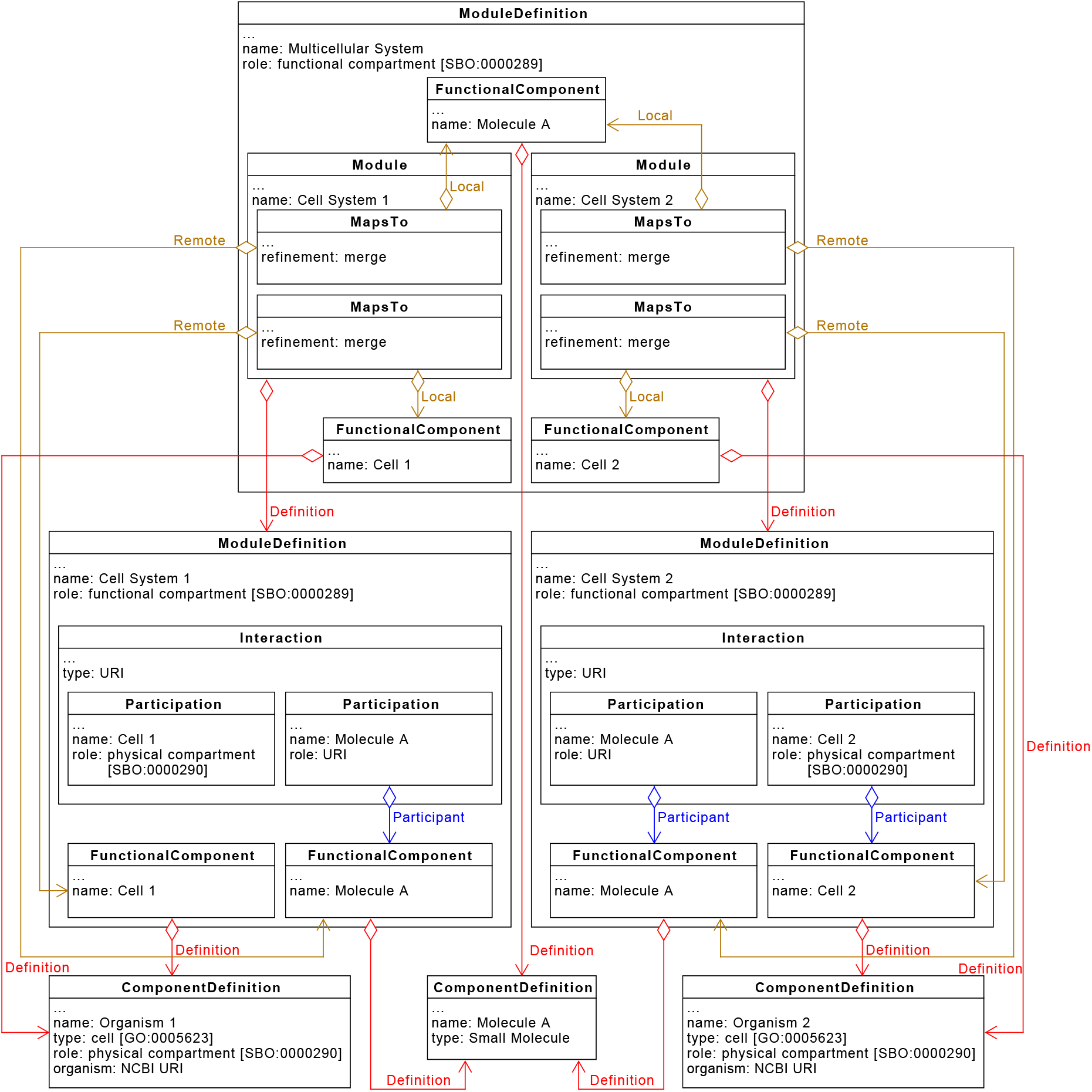
UML Diagram Depicting the Third Suggestion for Representing Multicellular Designs in SBOL: Captured here is a design involving two cells which both interact with the small molecule ‘Molecule A’. Designs for Cell 1 and Cell are captured using the approach depicted in *Figure 1B*, as this proposed approach is not compatible with the method shown in *Figure 1A*. The overall multicellular system is represented by a *ModuleDefinition* with a role of ‘functional compartment’, which is an SBO term. The two systems involving Cell 1 and Cell 2 are included in this multicellular design as instances of the *Module* class. Entities in the multicellular design, including those in the Cell 1 and Cell 2 system designs, are instantiated as a *FunctionalComponent*. Instances of the *MapsTo* class are used to map entities specified in both the individual cell systems, and the multicellular system.

### 2.5 Representing Composition of Multicellular Systems in SBOL

The proportion of cell types present in a multicellular system has been captured through the use of the ‘annotation’ property on classes which represent cells in the multicellular design. As a best practice, the value of these annotations is a number less than or equal to 1.0. These values are the fraction of a cell type present in the system compared to all other cell types present. Therefore, the sum of all these values specified in the system should be equal to 1.0. Alternatively, an instance of the *Measure* class from the newly accepted SBOL Enhancement Proposal (SEP) 028 (*24*) could be used to specify the percentage of each cell in the design. These approaches are shown in Figure 5.

**Figure 5:**
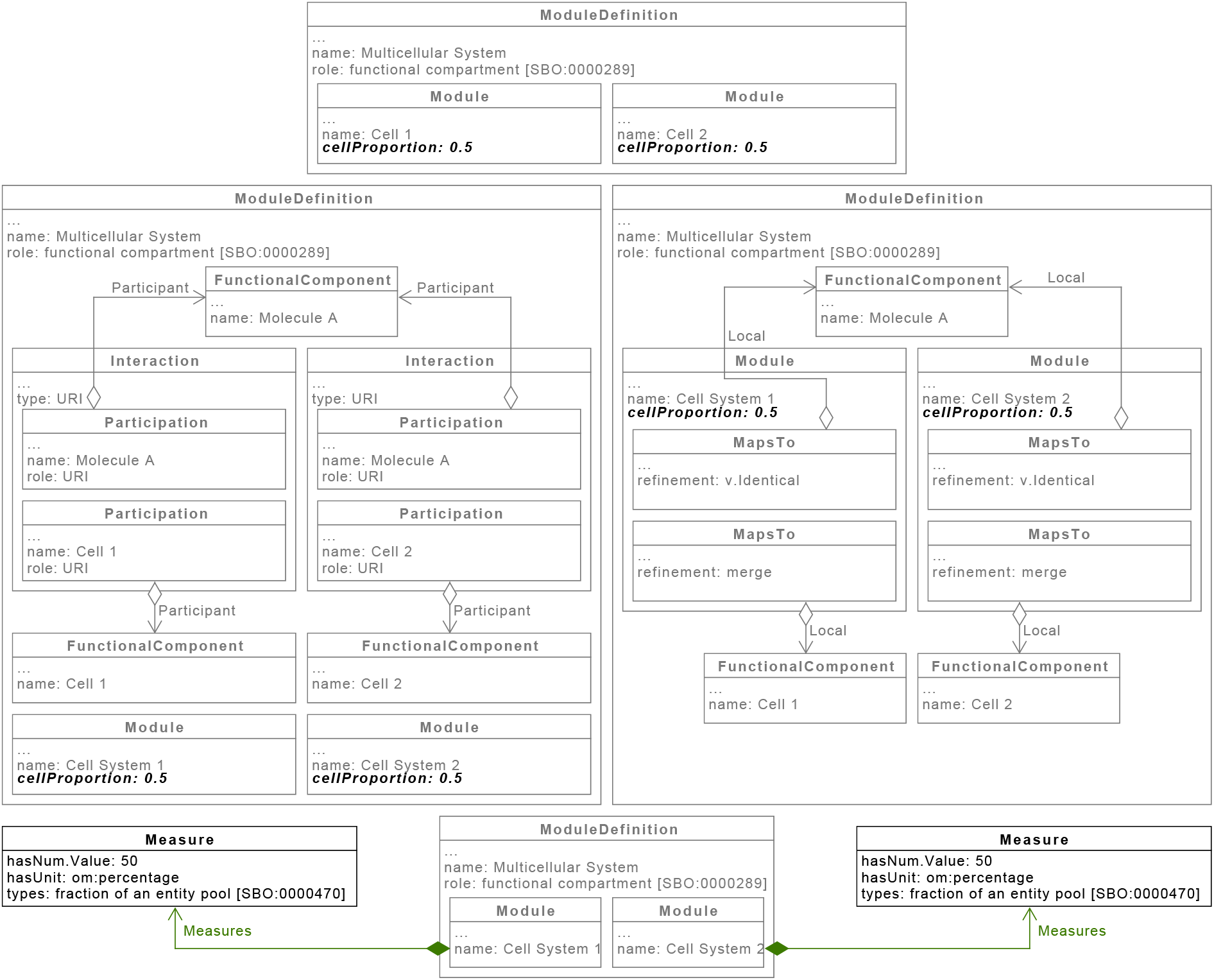
UML Diagram Depicting Suggestions for Capturing Cell Proportions in a Design: A-C) demonstrate which class instances should be annotated with cell proportions for the approaches described in Figures 2–4 respectively. D) Alternative to annotating class instances with cellular proportions based on SEP 028. Instances of the *Measure* class are used to capture the percentage of each cell type present in the multicellular system design.

## 3 Results and discussion

### 3.1 Example System: A Modular, Multicellular Biosensor

The Modular, Multicellular Biosensor (MMB) described by Newcastle iGEM 2017 (*25*) (Figure 6) is an example of a multicellular system. The MMB consists of three cell types: (i) a detector cell which converts the presence or absence of a specific stimulus into a genetic signal; (ii) a processor cell which modifies the signal from the detector cell in some way (e.g. amplifies it); and (iii) a reporter cell which converts the genetic signal into a response, such as a colour change or regulation of a metabolic pathway. The three cells have unidirectional communication, in which the detector cell passes a signal to the processor cell, and the processor cell passes a signal to the reporter cell. This communication is enabled using two quorum sensing (QS) mechanisms. The LasIR QS mechanism is used to pass the signal from the detector cell to the processor cell. When the stimulus is present the detector cell produces the acylhomoserine lactone (AHL) C12-HSL, which diffuses out of the detector cell and activates gene expression in the processor cell. The RhlIR QS mechanism is used to pass the signal from the processor to the reporter cell in a similar way, except the processor cell produces the AHL C4-HSL which activates gene expression in the reporter cell (*26*).

**Figure 6:**
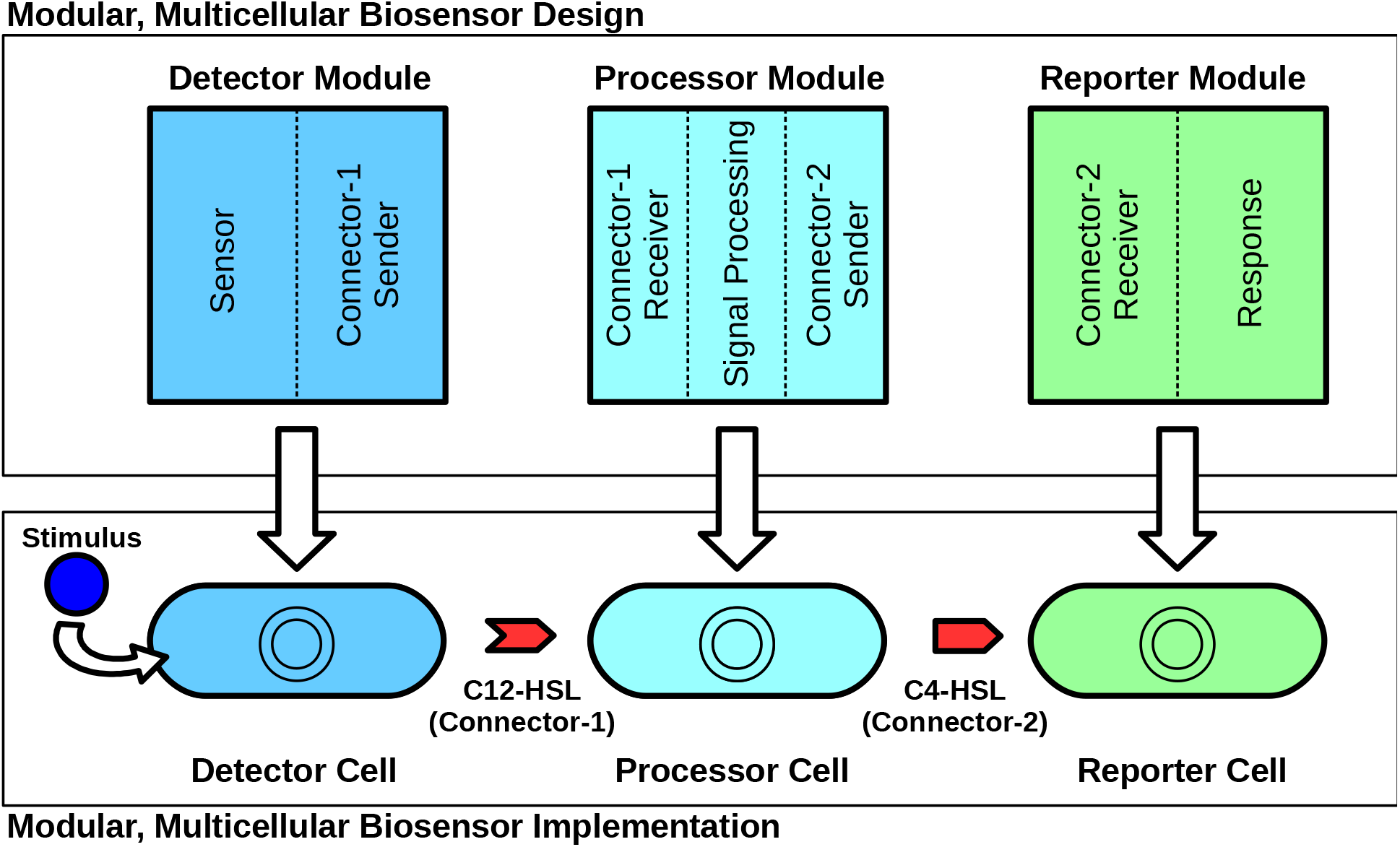
Schematic of a Generic Modular, Multicellular Biosensor. The Modular, Multicellular Biosensor Framework described by Newcastle iGEM 2017 consists of three modules, a detector, a signal processor, and a reporter. These three modules are expressed on separate plasmids and transformed in *E. coli* cells. A co-culture of these three cell types is then created to form a functional biosensor. The signal propagates from Detector cells to Processor cells to Reporter cells by use of AHL-based quorum sensing mechanisms.

The following sections discuss how each aspect of multicellular system designs can be represented in a standard way using SBOL. The MMB described above has been used as an example system to demonstrate the applicability of this approach.

### 3.2 A Standardised Representation of Cells

When attempting to re-produce a previously designed multicellular system, it is necessary to know some fundamental information about the cells used. The most crucial information is the strains used, any plasmids transformed into the cells, and expected functionality. By providing precise taxonomic information, it can be ensured that the correct strains are used when other researchers attempt to re-create a system, hence increasing reproducibility. It is suggested here that the organism’s species and strain be defined by providing a link to the relevant entry in the NCBI taxonomy database. This standardised approach would allow for easier automated retrieval of information about the organism. Whilst a link to an NCBI entry would be preferable, there are instances where this may not be possible (e.g. novel species not yet recorded). In these cases, it is suggested that instead a different database which does contain the organism is used. If the organism is not included in any databases, then a description of the organism should be provided instead. Including detailed information about any plasmids used in the cells is also essential, as plasmids will likely encode genetic circuits/expression devices which have a major effect on the cell’s behaviour. Finally, it is useful to record and share the exact functionality of one’s design to ensure that future users select the correct system for their desired application, and to allow for informed modification of the system.

Here, two approaches are proposed for representing the information stated above in a standardised format using SBOL. The main difference between the two approaches is that in the first, all of this information is captured in a single class instance, whereas in the second this information is split. These approaches are explained in detail here, using the IPTG Detector Cell in the Modular, Multicellular Biosensor (MMB) as an example.

The first approach (Figure 7A) uses an instance of the *ModuleDefinition* class to represent the cell. This class instance is annotated with a link to the NCBI taxonomy database to confer that it is an *E. coli* DH5 strain. Instances of the *FunctionalComponent* class are used to capture important molecules which are present within the cell. In this case, these molecules are the IPTG Detector plasmid DNA, IPTG, and C12 homoserine lactone (C12- HSL). An instance of the *Interaction* class is used to capture the functionality of the IPTG Detector Cell; IPTG induces a genetic construct encoded by the IPTG Detector plasmid to produce C12-HSL. The *FunctionalComponent* which represents the plasmid DNA is defined by an instance of the *ComponentDefinition* class, which stores structural information such as the sequence of the plasmid, and the names of genetic parts used.

**Figure 7:**
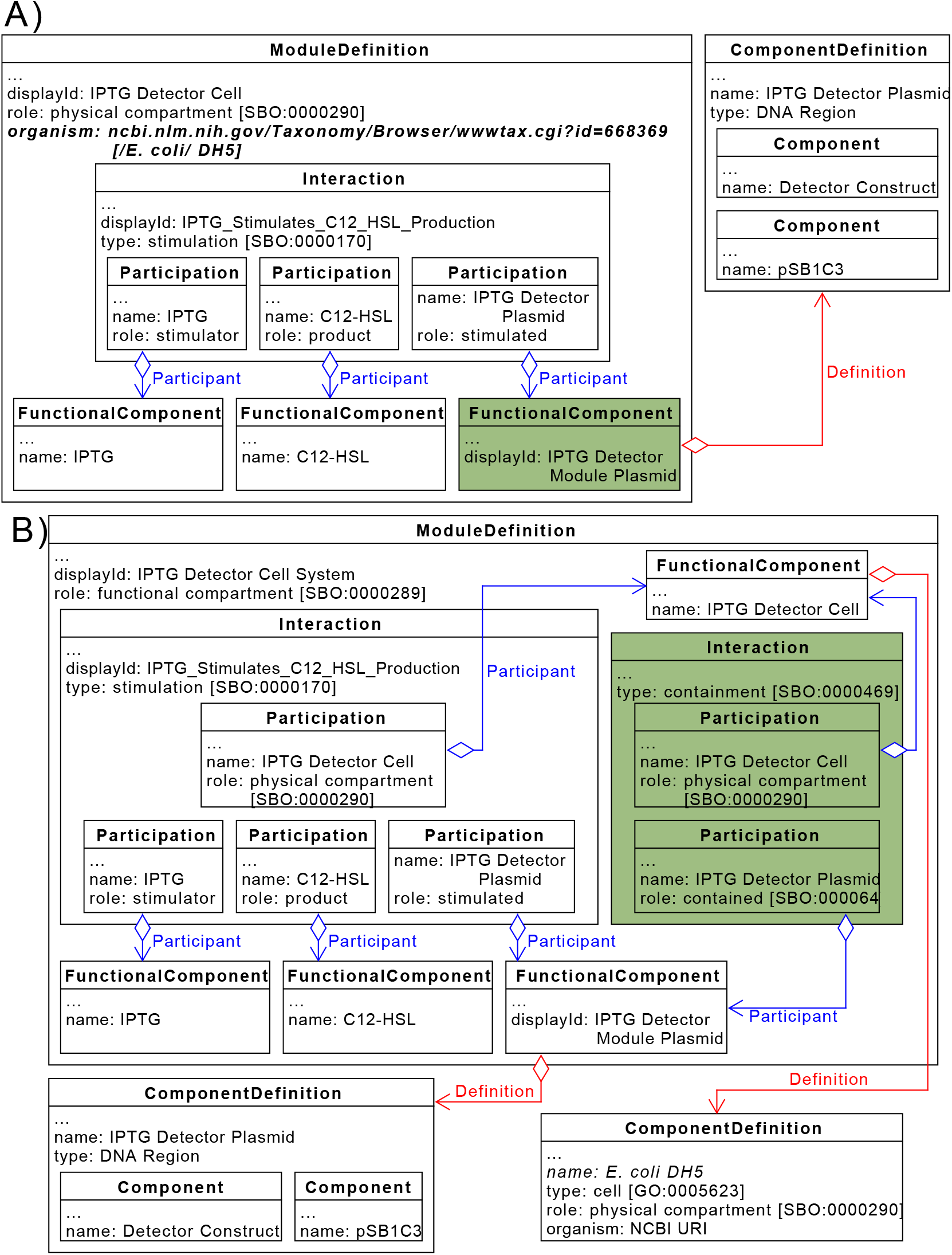
Capturing the IPTG Detector Cell Design from the Modular, Multicellular Biosensor using SBOL: Capturing the IPTG Detector Cell design using approach 1 depicted in Figure 1A (**A)** and approach 2 depicted in Figure 1B (**B)**. The cell strain used in the design is *E. coli* DH5α, which is entry 668369 in the NCBI Taxonomy Database. This cell contains the IPTG Detector Module Plasmid, which allows the cell to produce the small molecule C12-HSL (homoserine lactone) in the presence of the small molecule IPTG.

The second approach (Figure 7B) uses the *ModuleDefinition* instance to represent a system in which the cell is implemented, as opposed to the actual cell as in the first approach. Instead, a separate *ComponentDefinition* instance is used to capture taxonomic information about the cell, i.e. that it is an *E. coli* DH5 strain. This *ComponentDefinition* is used to define an instance of the *FunctionalComponent* class, which represents the cell within the system. As with the first approach, the important molecules in the design are also captured using instances of the *FunctionalComponent* class, and transformation of the cell with a plasmid can be captured in the same way. However, the containment of the plasmid within the cell could also be captured explicitly, using a ‘containment’ interaction which has the host cell and plasmid DNA as participants with roles of ‘physical compartment’ and ‘contained’ respectively. Other molecules which are produced by the cell and are not transported out into the extracellular space can also be defined in this way, if this information is important to the system’s design/function.

Both approaches capture the same information, but in different ways. The first approach is less complex, and captures information such as containment of molecules in an implicit way. However, it is possible to explicitly capture the movement of molecules in/out of the cell if desired by using the *Interaction* class to define specific transport mechanisms. The second approach is more complex, but captures information such as molecule containment explicitly. Additionally, the use of a separate *ComponentDefinition* class instance to capture taxonomic information make it easier to capture how a certain strain was derived from another strain without having to re-define all molecules and interactions occurring within the cell.

#### 3.2.1 A Standard Representation of Multicellular Systems

Once cells have been individually defined, they can be included in a multicellular system design. In systems involving more than one cell, it is important to capture the relative amounts of each cell type, as this can have a large effect on the system’s behaviour. Additionally, it should be possible to define how each cell type interacts with other cells in the system, as these intercellular interactions are usually the basis for a multicellular system’s functionality. Intercellular interactions normally occur by the same type of molecule being involved in processes of different cells. For example, two cell types in the system may require the same molecule for metabolic pathways to facilitate cell growth and hence are competing for resources, or one cell may produce a molecule which interacts with genetic circuits in a second cell, which is the basis for intercellular communication.

Proposed here are three approaches for implementing multiple cells into a single design. For all approaches, the multicellular system is represented by an instance of the *ModuleDefinition* class. The first approach involves simply instantiating each cell within this *ModuleDefinition* using the *Module* class. This approach is compatible with any method of representing cells discussed above. Intercellular interactions are captured implicitly, where interactions are elucidated by determining if any molecules are instantiated as a *FunctionalComponent* in more than one cell present in the multicellular system design. The type of intercellular interaction can be determined by comparing the molecule is doing in each cell. For example, if a metabolite is used in a metabolic process in two cells, then it can be determined that the cells are competing for resources. Another example is if one cell produces a molecule which is used by another cell to produce a product of interest. This would be an example of co-metabolism, where cells are co-operating to produce a desired product. Figure 8 shows how this approach could be used to capture intercellular communication between the IPTG Detector Cell and the Blank Processor Cell in the MMB. The IPTG Detector Cell produced C12-HSL in the presence of IPTG, and the Blank Processor Cell is stimulated by the C12-HSL to produce C4-HSL. Therefore, it can be determined that uni-directional communication from the Detector Cell to the Processor Cell is occurring, using C12-HSL.

**Figure 8:**
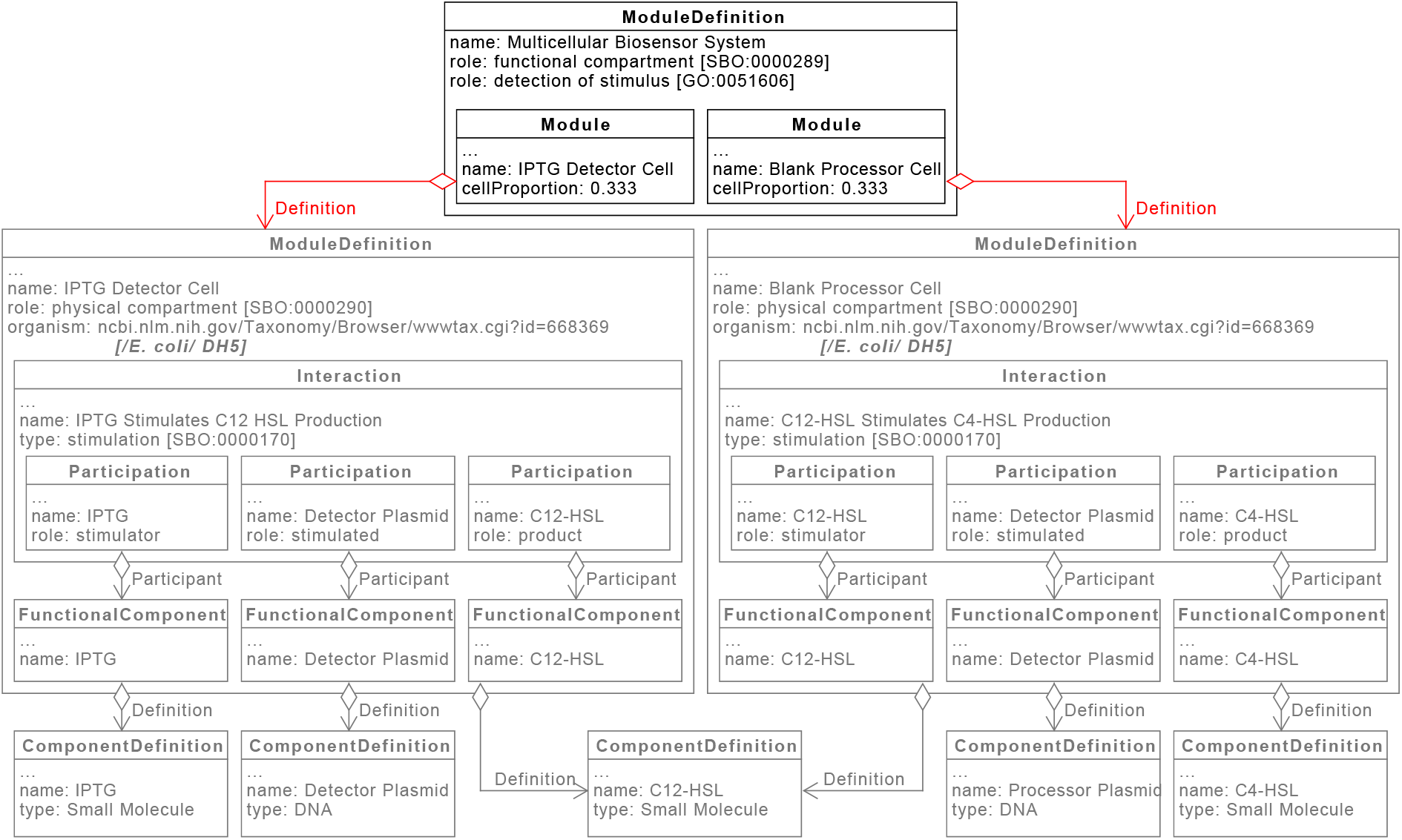
Capturing Interactions between the IPTG Detector and Blank Processor Cells using SBOL - Approach 1: Interactions between two cells in the Modular, Multicellular Biosensor are captured using the approach described in Figure 2. The Detector Cell produces C12-HSL in the presence of IPTG, and the Processor Cell produces C4-HSL in the presence of C12-HSL. This approach explicitly captures that the C12-HSL used in both cell designs is the same molecule as both instances are defined by the same *ComponentDefinition*. The cell/cell system designs are included in the same multicellular system design as instances of the *Module* class, and therefore it can be deduced that the C12-HSL produced by the Detector Cell will be available to the Processor Cell, allowing it to produce C4-HSL. This demonstrates unidirectional communication from the Detector Cell to the Processor Cell, using C12-HSL as the communication molecule.

The second approach explicitly captures interactions between cells. All cells and important molecules are included in the *ModuleDefinition* as instances of the *FunctionalComponent* class. The *Interaction* class is then used to define relevant process, using the *FunctionalComponent* representing the relevant cell as a participant with a role of ‘physical compartment’ to convey that the process in question is occurring within that cell. Interactions between cells are then determined by comparing interactions and seeing if any use the same *FunctionalComponent* as a participant. An example of this is shown in Figure 9, where this approach is used to once again to capture communication between the IPTG Detector Cell and the Processor Cell of the MMB.

**Figure 9:**
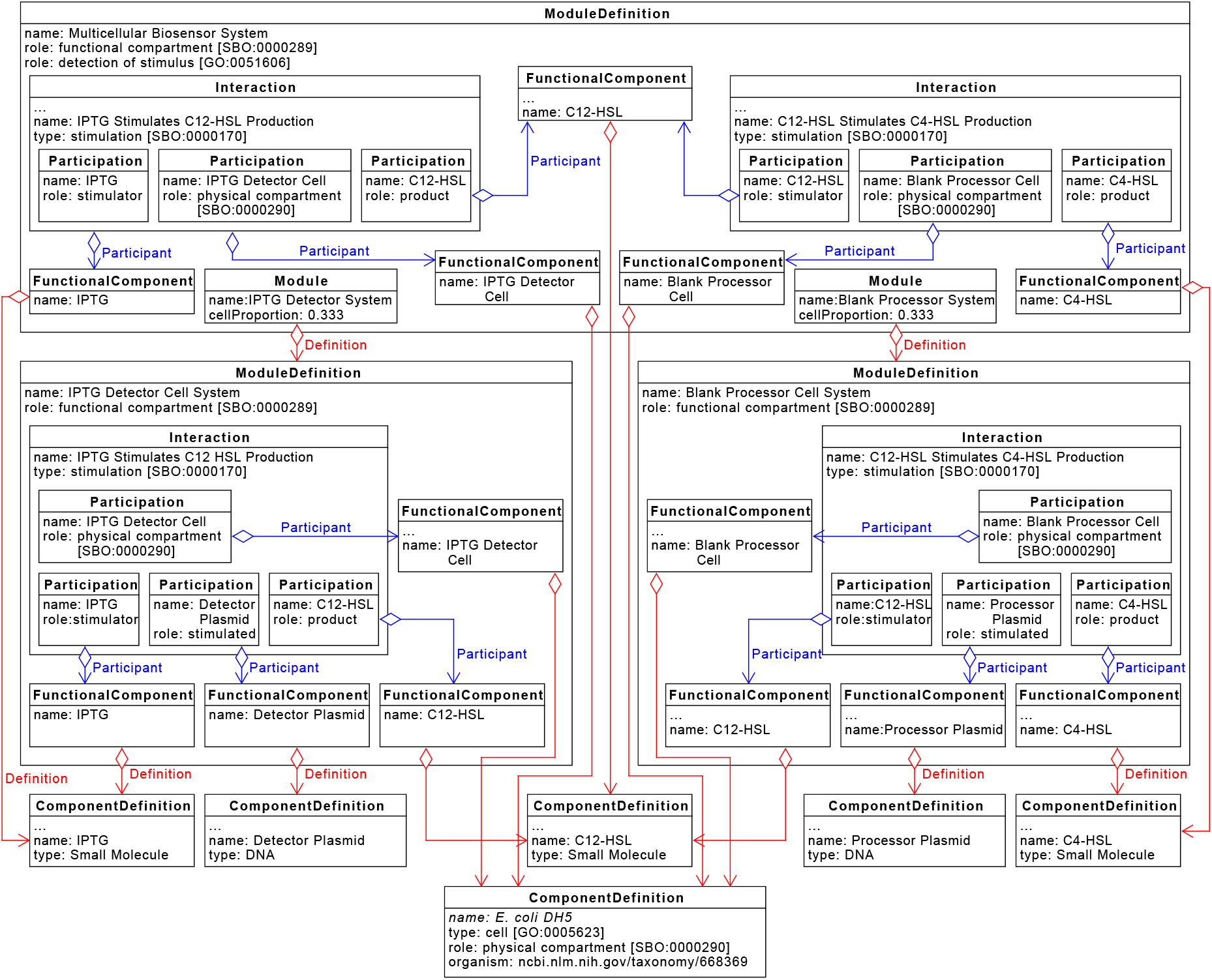
Capturing Interactions between the IPTG Detector and Blank Processor Cells using SBOL - Approach 2: Interactions between two cells in the Modular, Multicellular Biosensor are captured using the approach described in Figure 3. The Detector Cell produces C12-HSL in the presence of IPTG, and the Processor Cell produces C4-HSL in the presence of C12-HSL. This approach explicitly captures that the C12-HSL used in both cell designs is the same molecule as both instances are defined by the same *ComponentDefinition*. Two instances of the *E. coli* strains are included in the same multicellular system design as a *FunctionalComponent*, and labelled as an IPTG Detector Cell and Blank Processor Cell. Additionally, all important entities are also included as instances of the *FunctionalComponent* class. Two instances of the *Interaction* class are used to capture that the IPTG Detector Cell produced C12-HSL in the presence of IPTG, and that the Processor Cell produces C4-HSL in the presence of C12-HSL. As both interactions use the same *FunctionalComponent* representing the C12-HSL, it can be determined that the cells particpate in unidirectional communication using C12-HSL as a signalling molecule.

Finally, the third approach proposed here is a combination of both previous suggestions. It should be noted here that this approach would only be compatible with the second method of representing cells in SBOL described above. As with the first approach, the cell systems are included in the multicellular design by instantiating them as a *Module*, and as with the second important molecules as included as a *FunctionalComponent*. Additionally, the cells themselves are included as a *FunctionalComponent*, which is defined by the *ComponentDefinition* used to capture the taxonomic information about the organism. The *Module* instances contain *MapsTo* class instances, which are used to explicitly capture that entities in the lower level cell systems are the same as those used in the multicellular system design, as opposed to determining this implicitly. An example of how this approach can be used to capture communication from the IPTG Detector Cell to the Processor Cell in the MMB is shown in Figure 10.

**Figure 10:**
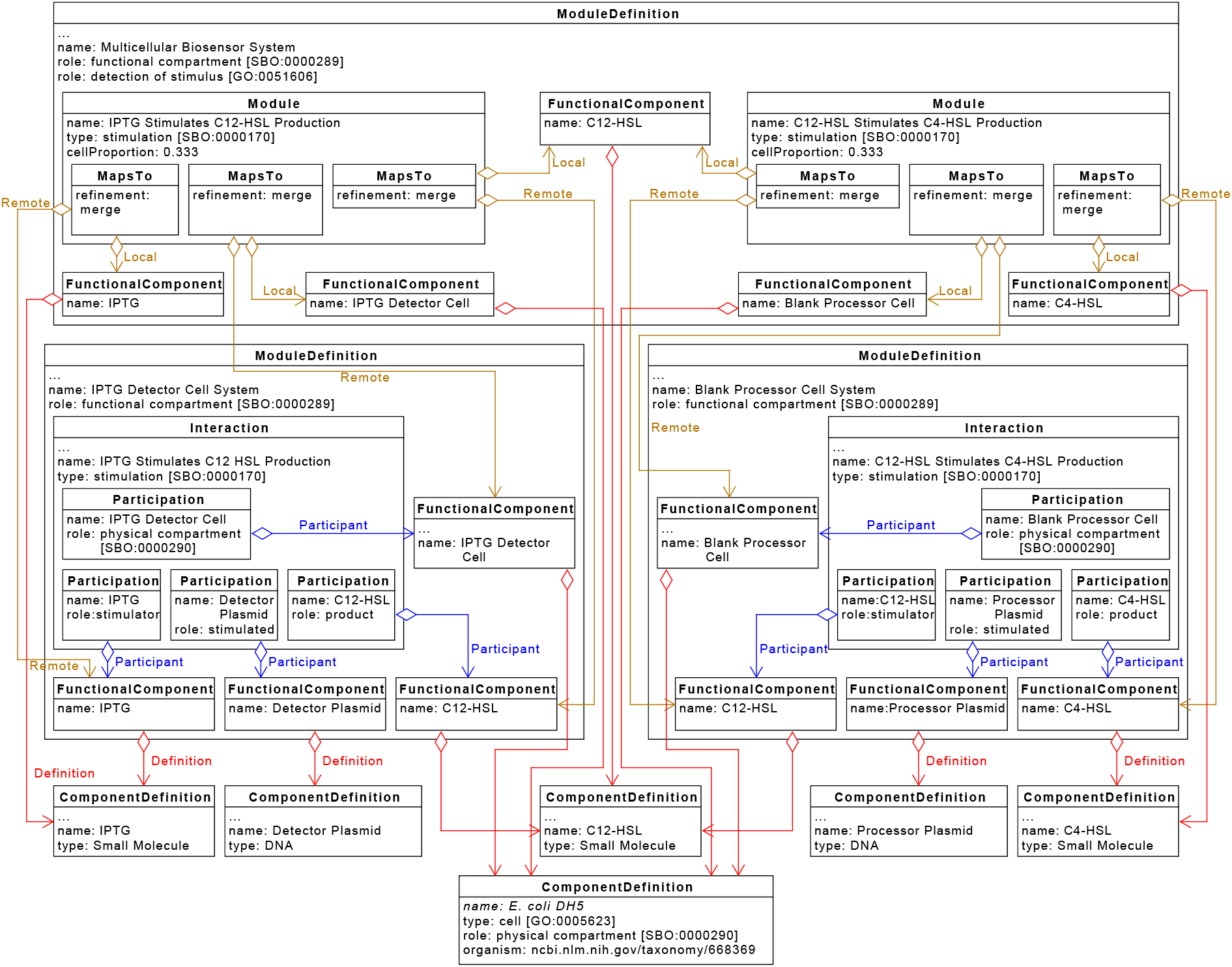
Capturing Interactions between the IPTG Detector and Blank Processor Cells using SBOL - Approach 3: Interactions between two cells in the Modular, Multicellular Biosensor are captured using the approach described in Figure 3. The Detector Cell produces C12-HSL in the presence of IPTG, and the Processor Cell produces C4-HSL in the presence of C12-HSL. This approach explicitly captures that the C12-HSL used in both cell designs is the same molecule as both instances are defined by the same *ComponentDefinition*. Two instances of the *E. coli* strains are included in the same multicellular system design as a *FunctionalComponent*, and labelled as an IPTG Detector Cell and Blank Processor Cell. Additionally, all important entities are also included as instances of the *FunctionalComponent* class. The two system designs involving the Detector and Processor Cells are also instantiated as a *Module*. Instances of the *MapsTo* class are used to link identical entities in both the multicellular and unicellular designs. For example, the IPTG small molecules in the multicellular design is linked to the IPTG molecule in the Detector Cell system design via the *MapsTo* class, indicating that they are the same molecule.

As explained above, it is also important to capture the relative amounts of each cell type present in the multicellular system. It is proposed that the classes used to represent cells in the multicellular system are annotated with the fractional amount of the total cell population each specific cell occupies. For example, if the Detector, Processor, and Reporter cells are present in a 1:1:1 ratio, then each cell would be annotated with a fractional amount of 0.33. This is shown in Figures 8, 9, and 10.

Each approach described has advantages and disadvantages. The first approach is less complicated, however it may be more difficult in some designs to determine if cells are interacting. The second approach avoids this issue by requiring designers to explicitly include intercellular interactions at the level of the multicellular system. However, this approach can lead to duplication of interactions in the design, where the same interaction is specified at both the level of the cell’s design, and the multicellular system design. The third approach allows important molecules to be defined at the level of the multicellular system without duplicating interactions, though the specific multicellular-level interactions are still defined implicitly. Additionally, this approach enables one to map between different pools of the same molecule, which is useful when some pools are accessible extracellularly but others are not. This capability comes, however, at the cost of greater complexity than the previous two approaches.

Despite the issues mentioned above, defining interactions between cells implicitly could be beneficial. Automated design software could combine cell designs into a multicellular system and automatically determine the nature of any interactions between cells. However, if designs for cells are obtained from other researchers, perhaps via a database such as SynBioHub (*27*), processes which are not important in a homogeneous design but are crucial in a multicellular system may be missing, and hence important interactions lost. An example of this would be metabolism of a certain molecule. In a homogeneous culture this may not be important, however if this cell is used in a multicellular system, and another cell is present which also uses the same metabolite, then competition between the two cells will occur, however this information will not be present in the design.

### 3.3 Representing Multicellular System in Future Versions of SBOL

As explained previously, the proposed approach for capturing information about a cell’s design described in Figure 1A is not compatible with the approaches for including multiple cells in a single design described in Figures 3 and 4. This is because the cell designs are captured using instances of the *ModuleDefinition* class, and the approaches in Figures 3 and 4 require cells to be instantiated as a *FunctionalComponent*. According to the SBOL version 2 specifications, instances of the *FunctionalComponent* class cannot have a definition which points to a *ModuleDefinition*. However, a proposal has been put forward for combining the *ModuleDefinition* and *ComponentDefinition* classes in a future version of SBOL, where a *ComponentDefinition* can contain *Interaction* and *FunctionalComponent* class instances (*28*). Additionally, the *FunctionalComponent* class would also be merged with the *Component* class. In this case, the cell designs can be captured using a *ComponentDefinition*, and therefore be instantiated in a multicellular system design as a *Component*. An example of this approach is shown in Figure 11, where the IPTG Detector Cell and Blank Processor Cell designs are captured using the approach in Figure 1A, and included in a multicellular system design using the approach described in Figure 3.

**Figure 11:**
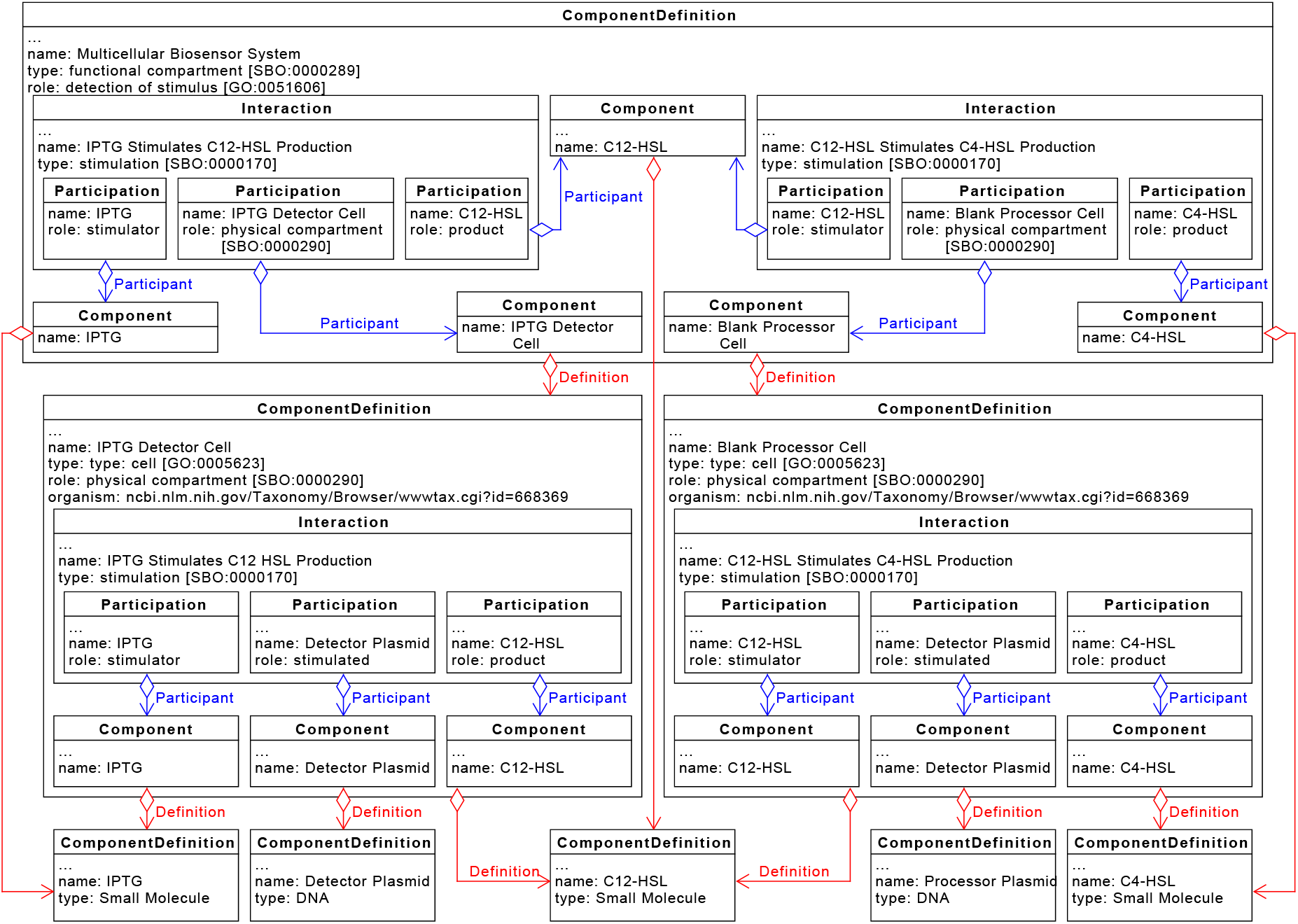
Capturing Interactions between the IPTG Detector and Blank Processor Cells using a Future Version of SBOL: The IPTG Detector and Blank Processor Cells are represented using instances of the *ComponentDefinition* class in a theoretical future version of SBOL where the *ModuleDefinition* class has been merged with the *ComponentDefinition* class. The designs of each cell are captured as described in Figure 7A. These cell designs are both included in a multicellular system design which is as instances of the *Component* class. Interactions between cells are defined in the same way as the approach in Figure 9.

## 4 Conclusion

Whilst the SBOL data model has been used in the past to capture information about genetic constructs and intracellular interactions, it has not been widely used to describe and share information about multicellular systems. This paper aimed to illustrate how multicellular system designs can be captured in a standard way using SBOL. Examples have been provided to help illustrate specific concepts and demonstrate the feasibility of each approach described. It is hoped that the approaches described here can be used to facilitate discussion on how SBOL can be used/modified to enable easy sharing of multicellular designs. This should pave the way for the development of software tools which can aid researchers in designing and sharing synthetic multicellular systems.

## Acknowledgement

This document does not contain technology or technical data controlled under either the U.S. International Traffic in Arms Regulations or the U.S. Export Administration Regulations.

